## Supplemental Material for "Mechanistic computational modeling of sFLT1 secretion dynamics"

This document contains the following supplemental material related to the manuscript "Mechanistic computational modeling of sFLT1 secretion dynamics":

- Supplemental Methods
- Supplemental Results
- Supplemental Tables S1-S3
- Supplemental Figures S1-S13
- Supplemental References

### Supplemental Methods

Here we derive additional equations and relationships used in analysis of these ODE and DDE models of sFLT1 secretion. Some equations are repeated from the main text for clarity, and are numbered as in the main text.

**Ordinary differential equations (ODEs).** The rate of change of intracellular sFLT1 ($\frac{dI(t)}{dt}$) is the balance of production, secretion, and intracellular degradation:

$$\frac{dI(t)}{dt} = \alpha - \beta \cdot I(t) - \gamma \cdot I(t) \qquad \text{(Eq. 1)}$$

The rate of change of extracellular sFLT1 ($\frac{dX(t)}{dt}$) is the balance of secretion and extracellular degradation:

$$\frac{dX(t)}{dt} = \beta \cdot I(t) - \delta \cdot X(t) \qquad \text{(Eq. 2)}$$

Units of *I*, *X*: #/cell (number of proteins per cell)

Units of α: #/cell/h (number of proteins per cell per hour)

Units of β, γ, δ: 1/h (inverse hours)

**Delay differential equations (DDEs).** These equations apply a time delay, $\tau$, to intracellular sFLT1 when calculating secretion rate and intracellular degradation rate:

$$\frac{dI(t)}{dt} = \alpha - \beta \cdot I(t-\tau) - \gamma \cdot I(t-\tau) \qquad \text{(Eq. 3)}$$

$$\frac{dX(t)}{dt} = \beta \cdot I(t-\tau) - \delta \cdot X(t) \quad \text{(Eq. 4)}$$

Units of $\tau$: h (hours)

**Observed constraints.** Optimization of our model to experimental data yielded the following constants:

$$c_1 = \alpha\beta$$

i.e., the product of production rate and secretion rate constant is constrained to a single value.

$$c_2 = \beta + \gamma$$

i.e., the sum of secretion and intracellular degradation rate constants is constrained to a single value.

**Theoretical steady state $(I_{SS}, X_{SS})$ of the ODE and DDE systems.** At steady state: $\frac{dI(t)}{dt} = \frac{dX(t)}{dt} = 0$, $I(t) = I_{SS}$, and $X(t) = X_{SS}$.

From Eq. 1:

$$0 = \alpha - (\beta + \gamma) \cdot I_{SS} \Rightarrow \alpha = (\beta + \gamma) \cdot I_{SS} \Rightarrow I_{SS} = \frac{\alpha}{\beta+\gamma} \quad \text{(Eq. 6)}$$

From Eq. 2:

$$0 = \beta \cdot I_{SS} - \delta \cdot X_{SS} \Rightarrow \beta \cdot I_{SS} = \delta \cdot X_{SS} \Rightarrow X_{SS} = \frac{\alpha\beta}{\delta(\beta+\gamma)} \quad \text{(Eq. 7)}$$

In terms of the observed constraints $c_1 = \alpha\beta$ and $c_2 = \beta + \gamma$:

$$I_{SS} = \frac{\alpha}{\beta+\gamma} = \frac{\alpha}{c_2} = \frac{c_1}{\beta c_2} \quad \text{(Eq. 6a)}$$

$$X_{SS} = \frac{\alpha\beta}{\delta(\beta+\gamma)} = \frac{c_1}{\delta c_2} = I_{SS}\frac{\beta}{\delta} \quad \text{(Eq. 7a)}$$

For the DDE system, $I(t-\tau) = I(t) = I_{SS}$, and $X(t-\tau) = X(t) = X_{SS}$. Thus, we get the same results as Eq. 6a and Eq. 7a.

**Characteristic time to half-steady-state for intracellular sFLT1 ($T_{50\_I}$) for the ODE system.** Assuming first order secretion rate constant β and first order intracellular degradation rate constant γ, *I* has a combined first order elimination rate constant $\beta + \gamma$. From first principles of kinetics, accumulation of *I* to $I_{SS}$ is:

$$I(t) = I_{SS} \cdot (1 - e^{-(\beta+\gamma)\cdot t})$$

At $t = T_{50\_I}$, when $I(T_{50\_I}) = \frac{I_{SS}}{2}$:

$$\frac{I_{SS}}{2} = I_{SS} \cdot (1 - e^{-(\beta+\gamma)\cdot T_{50\_I}}) \Rightarrow \frac{1}{2} = 1 - e^{-(\beta+\gamma)\cdot T_{50\_I}} \Rightarrow e^{-(\beta+\gamma)\cdot T_{50\_I}} = \frac{1}{2} \Rightarrow$$

$$-(\beta + \gamma) \cdot T_{50\_I} = ln\ \frac{1}{2} \Rightarrow$$

$$T_{50\_I} = \frac{ln\ 2}{\beta+\gamma} \qquad \text{(Eq. 8)}$$

**Characteristic time to half-steady-state for extracellular sFLT1 ($T_{50\_X}$) for the ODE system.** From first principles of kinetics, assuming first order extracellular degradation rate constant δ accumulation of *X* to $X_{SS}$ is:

$$X(t) = X_{SS} \cdot (1 - e^{-\delta\cdot t})$$

At $t = T_{50\_X}$, when $I(T_{50\_X}) = \frac{X_{SS}}{2}$:

$$\frac{X_{SS}}{2} = X_{SS} \cdot (1 - e^{-\delta\cdot T_{50\_X}}) \Rightarrow \frac{1}{2} = 1 - e^{-\delta\cdot T_{50\_X}} \Rightarrow e^{-\delta T_{50\_X}} = \frac{1}{2} \Rightarrow$$

$$-\delta \cdot T_{50\_I} = ln\ \frac{1}{2} \Rightarrow$$

$$T_{50\_X} = \frac{ln\ 2}{\delta} \qquad \text{(Eq. 9)}$$

**Process fluxes.** These non-negative fluxes represent the rates over time of each modeled process in #/cell/h, and are the constituent rate terms from the right hand side of the DDE equations given earlier. The fluxes for the ODEs are identical for $\tau = 0$. Prod = production, Secr = secretion, IDeg = intracellular degradation, XDeg = extracellular degradation.

$$\Phi_{Prod}(t) = \alpha$$
$$\Phi_{Secr}(t) = \beta \cdot I(t-\tau)$$
$$\Phi_{IDeg}(t) = \gamma \cdot I(t-\tau)$$
$$\Phi_{XDeg}(t) = \delta \cdot X(t)$$

**Steady state secretion flux** ($\Phi_{Secr}$)**.** At steady state, from Eq. 6a, $I(t-\tau) = I_{SS} = \frac{\alpha}{\beta+\gamma}$.

Therefore:

$$\Phi_{Secr} = \beta \cdot I_{SS} = \frac{\alpha\beta}{\beta+\gamma} = \frac{c_1}{c_2}$$

**Steady state relationship between intracellular degradation flux** ($\Phi_{IDeg}$)**and production flux** ($\Phi_{Prod}$)**.** At steady state, from Eq. 6a, $I(t-\tau) = I_{SS} = \frac{\alpha}{c_2} = \frac{\Phi_{Prod}}{c_2}$. Therefore:

$$\Phi_{IDeg} = \gamma \cdot \frac{\Phi_{Prod}}{c_2} = \frac{c_2-\beta}{c_2} \cdot \Phi_{Prod} = (1-\frac{\beta}{c_2}) \cdot \Phi_{Prod} = \Phi_{Prod} - \frac{\alpha\beta}{c_2} \Rightarrow$$
$$\Phi_{IDeg} = \Phi_{Prod} - \frac{c_1}{c_2}$$

**Asymptotic behavior of steady state intracellular degradation flux** ($\Phi_{IDeg}$)**.** Since $\Phi_{IDeg}$ is non-negative, $min(\Phi_{IDeg}) = 0$, and $\Phi_{Prod} \geq \frac{c_1}{c_2}$.

When $\Phi_{Prod} \gg \frac{c_1}{c_2}$, $\Phi_{IDeg} = \Phi_{Prod} - \frac{c_1}{c_2} \rightarrow \Phi_{Prod}$.

i.e. maximum intracellular degradation is equal to production, and this occurs when the secretion rate constant is zero ($\beta \rightarrow 0$).

**Theoretical lower bound of production flux ($\Phi_{Prod}$).** $\Phi_{Prod}$ increases monotonically with $\Phi_{IDeg}$, so $min(\Phi_{Prod})$ occurs when $\Phi_{IDeg} = min(\Phi_{IDeg})$. Therefore:

$$min(\Phi_{Prod}) = min(\Phi_{IDeg}) + \frac{c_1}{c_2} = 0 + \frac{c_1}{c_2} = \frac{c_1}{c_2}$$

i.e. production is minimized when there is no intracellular degradation, only secretion.

**Steady state relationship between intracellular sFLT1 ($I_{SS}$) and intracellular degradation flux ($\Phi_{IDeg}$).**

$$I_{SS} = \frac{\Phi_{Prod}}{c_2} \Rightarrow \Phi_{Prod} = c_2 I_{SS}$$

$$\Phi_{IDeg} = \Phi_{Prod} - \frac{c_1}{c_2} = c_2 I_{SS} - \frac{c_1}{c_2} \Rightarrow c_2 I_{SS} = \Phi_{IDeg} + \frac{c_1}{c_2} \Rightarrow$$

$$I_{SS} = \frac{\Phi_{IDeg}}{c_2} + \frac{c_1}{{c_2}^2}$$

**Asymptotic behavior of steady state intracellular sFLT1 ($I_{SS}$).** Since both $I_{SS}$ and $\Phi_{Prod}$ are non-negative and directly proportional, $min(I_{SS})$ occurs when $\Phi_{Prod} = min(\Phi_{Prod})$. Therefore:

$$min(I_{SS}) = min(\Phi_{Prod})/c_2 = \frac{c_1/c_2}{c_2} \Rightarrow min(I_{SS}) = \frac{c_1}{{c_2}^2}$$

This is also consistent with $I_{SS} = \frac{\Phi_{IDeg}}{c_2} + \frac{c_1}{{c_2}^2}$ as $\Phi_{IDeg} \to 0$.

When $\Phi_{IDeg} \gg \frac{c_1}{c_2}$, $\Phi_{IDeg} + \frac{c_1}{c_2} \to \Phi_{IDeg}$ and $I_{SS} = \frac{1}{c_2}(\Phi_{IDeg} + \frac{c_1}{c_2}) \to \frac{\Phi_{IDeg}}{c_2}$.

### Supplemental Results

**General analysis of ODE model of secretion.** Analysis of model solution dynamics (**Figure S1A**) shows that in response to a step increase in the rate of production from 0 to $\alpha$ at time $t = 0$, intracellular sFLT1 increases hyperbolically to steady state, and extracellular sFLT1 increases to its steady state with a delay relative to pure first-order kinetics (which would result in $X$ reaching $X_{SS}/2$ at a normalized time of 1); this delay is due to the time needed for production and secretion to occur. The system has an equal but opposite response to a step decrease in production. Phase plane analysis confirms that intracellular, extracellular, and total sFLT1 increase monotonically when production increases and decrease monotonically when production decreases, and that intracellular sFLT1 approaches its steady state faster than extracellular sFLT1 (**Figure S1B**).

**General analysis of DDE model of secretion.** In an example solution to the base DDE model (**Figures S2A-B)**, intracellular sFLT1 initially responds to a step increase in production by overshooting its theoretical steady state value $I_{SS}$, while extracellular sFLT1 remains at 0 until $t = \tau$. As sFLT1 secretion begins, extracellular sFLT1 accumulates, while the intracellular sFLT1 first slows in accumulation rate and then decreases sharply below $I_{SS}$. Because of the delay, sFLT1 secretion initially continues to increase even as intracellular sFLT1 decreases, and extracellular sFLT1 also exceeds its theoretical steady state value $X_{SS}$. As intracellular sFLT1 falls, its production exceeds its clearance, and intracellular sFLT1 once again begins to accumulate and shows damped oscillations converging towards $I_{SS}$. These oscillations drive similar damped oscillations in extracellular sFLT1 converging towards $X_{SS}$, and because the oscillations are out of phase, total sFLT1 also oscillates rather than monotonically approaching steady state.

**Phase plane analysis of DDE model for varying secretion delay.** While the ODE model generated similar solution dynamics and phase plane portraits across a large region of parameter space, the DDE model solution shape varied significantly with the delay $\tau$, which changed both the amplitude and frequency of oscillations (**Figure S2C-D**). Some parameter regimes created dynamics that could not converge to steady state because they generated situations where intracellular sFLT1 clearance flux exceeded the available amount of intracellular sFLT1, a situation incompatible with conservation of mass. We therefore concluded that some parameter combinations were not physically permissible using the DDE model, and we excluded those combinations from future analysis.

**Sensitivity to values of constraints $c_1$ and $c_2$.** To better characterize the impact of observed constraints $(c_1, c_2)$, we varied one of these values while holding the other constant and simulated constitutive secretion. First, we allowed $\alpha$ and $\beta$ to vary while maintaining constant $c_1 = \alpha\beta = 7270$ #/cell/h$^2$, resulting in a range of values for $c_2 = \beta + \gamma$ . When $c_1$ is fixed, larger values of $c_2$ generate lower intracellular and extracellular sFLT1 levels (**Figure S10A**). Under these conditions, increasing $\alpha$ causes a linear increase in intracellular sFLT1 but a hyperbolic increase in extracellular sFLT1, while both intracellular and extracellular sFLT1 are inversely proportional to β and $c_2$ (**Figure S10B**).

Then, we allowed $(\beta, \gamma)$ to vary while maintaining constant $c_2 = \beta + \gamma = 0.173$ h$^{-1}$, resulting in a range of values for $c_1 = \alpha\beta$. When $c_2$ is fixed, intracellular sFLT1 remains constant independent of $c_1$, but extracellular sFLT1 increases as $c_1$increases (**Figure S10C**). The extracellular sFLT1 concentration increases linearly with increasing $c_1$ and $\beta$, and decreases linearly with increasing $\gamma$ (**Figure S10D**).

### Supplemental Tables

#### Supplemental Table S1

**Supplemental Table S1. Complete set of equations for candidate models.** Note that model A is equivalent to the "ODE-based model" and model B is equivalent to the "DDE-based model" described above.

| Model | ε | κ | τ | Equations |
|---|---|---|---|---|
| M1 | N | N | N | $\frac{dI(t)}{dt} = \alpha - \beta \cdot I(t) - \gamma \cdot I(t)$ <br> $\frac{dX(t)}{dt} = \beta \cdot I(t) - \delta \cdot X(t)$ |
| M2 | N | N | Y | $\frac{dI(t)}{dt} = \alpha - \beta \cdot I(t-\tau) - \gamma \cdot I(t-\tau)$ <br> $\frac{dX(t)}{dt} = \beta \cdot I(t-\tau) - \delta \cdot X(t)$ |
| M3 | N | Y | N | $\frac{dI(t)}{dt} = \alpha e^{-\kappa t} - \beta \cdot I(t) - \gamma \cdot I(t)$ <br> $\frac{dX(t)}{dt} = \beta \cdot I(t) - \delta \cdot X(t)$ |
| M4 | N | Y | Y | $\frac{dI(t)}{dt} = \alpha e^{-\kappa t} - \beta \cdot I(t-\tau) - \gamma \cdot I(t)$ <br> $\frac{dX(t)}{dt} = \beta \cdot I(t-\tau) - \delta \cdot X(t)$ |
| M5 | Y | N | N | $\frac{dI(t)}{dt} = \alpha - \beta \cdot I(t) - \gamma \cdot I(t) + \epsilon \cdot X(t)$ |

| | | | | |
|---|---|---|---|---|
| | | | | $\frac{dX(t)}{dt} = \beta \cdot I(t) - \delta \cdot X(t) - \epsilon \cdot X(t)$ |
| M6 | Y | N | Y | $\frac{dI(t)}{dt} = \alpha - \beta \cdot I(t-\tau) - \gamma \cdot I(t-\tau) + \epsilon \cdot X(t)$ <br> $\frac{dX(t)}{dt} = \beta \cdot I(t-\tau) - \delta \cdot X(t) - \epsilon \cdot X(t)$ |
| M7 | Y | Y | N | $\frac{dI(t)}{dt} = \alpha e^{-\kappa t} - \beta \cdot I(t) - \gamma \cdot I(t) + \epsilon \cdot X(t)$ <br> $\frac{dX(t)}{dt} = \beta \cdot I(t) - \delta \cdot X(t) - \epsilon \cdot X(t)$ |
| M8 | Y | Y | Y | $\frac{dI(t)}{dt} = \alpha e^{-\kappa t} - \beta \cdot I(t-\tau) - \gamma \cdot I(t) + \epsilon \cdot X(t)$ <br> $\frac{dX(t)}{dt} = \beta \cdot I(t-\tau) - \delta \cdot X(t) - \epsilon \cdot X(t)$ |

### Supplemental Table S2

**Supplemental Table S2. Characteristics of sFLT1 datasets used for mechanistic model optimization.**

Secretion scenario: C = constitutive, P = pulse-chase; Assay: E = ELISA, W = Western blot / autoradiography; Time points: * = flattened by normalization, † = omitted (see Methods for rationale).

| Dataset | Reference | Secretion scenario | Assay | X unit | X time points (h) | I unit | I time points (h) | # of data points ($AIC_C$) |
|---|---|---|---|---|---|---|---|---|
| Hornig | Hornig et al., 2000 | C | E | ng/mL | 0*, 3†, 6, 9, 12, 24, 48, 72 | NA | NA | 6 |
| Jung | Jung et al., 2012 | P | W | $\frac{X}{X_{8h}}$ | 0*, 2, 4, 6, 8*, 10† | $\frac{I}{I_{0h}}$ | 0*, 2, 4, 6, 8, 10 | 8 |
| Kinghorn | Kinghorn et al., 2024 | C | W | $\frac{X}{X_{24h}}$ | 0, 1, 2, 4, 8, 12, 24* | $\frac{I}{I_{0h}}$ | 0*, 1, 2, 4, 8, 12, 24 | 12 |

### Supplemental Table S3

**Supplemental Table S3.** Sampling distributions for initial parameter values and bounds for final parameter values during mechanistic model optimization.

| Parameter | Description | Distribution | Sampling bounds | | Optimization bounds | | Units |
|---|---|---|---|---|---|---|---|
| | | | Lower | Upper | Lower | Upper | |
| $\alpha$ | Production rate constant | Log uniform | $5 \times 10^{3}$ | $5 \times 10^{6}$ | ODE: $1 \times 10^{3}$ | ODE: $1 \times 10^{7}$ | #/cell/h |
| | | | | | other: $1 \times 10^{-4}$ | other: $1 \times 10^{7}$ | |
| $\beta$ | Secretion rate constant | Log uniform | $1 \times 10^{-3}$ | 1 | ODE: $1 \times 10^{-6}$ | ODE: 1 | 1/h |
| | | | | | other: $1 \times 10^{-4}$ | other: $1 \times 10^{7}$ | |
| $\gamma$ | Intracellular degradation rate constant | Log uniform | $1 \times 10^{-3}$ | 1 | ODE: $1 \times 10^{-6}$ | ODE: 1 | 1/h |
| | | | | | other: $1 \times 10^{-4}$ | other: $1 \times 10^{7}$ | |
| $\delta$ | Extracellular degradation rate constant | Log uniform | $1 \times 10^{-3}$ | 1 | ODE: $1 \times 10^{-6}$ | ODE: 1 | 1/h |
| | | | | | other: $1 \times 10^{-4}$ | other: $1 \times 10^{7}$ | |
| $\epsilon$ | Internalization rate constant | Log uniform | $1 \times 10^{-3}$ | 1 | $1 \times 10^{-4}$ | $1 \times 10^{7}$ | 1/h |
| $\kappa$ | Production decay [chase] constant | Uniform | 0 | 10 | $1 \times 10^{-4}$ | $1 \times 10^{7}$ | 1/h |
| $\tau$ | Maturation time delay | Uniform | 1 | 4 | $1 \times 10^{-4}$ | $1 \times 10^{7}$ | h |

### Supplemental Figures

#### Supplemental Figure S1

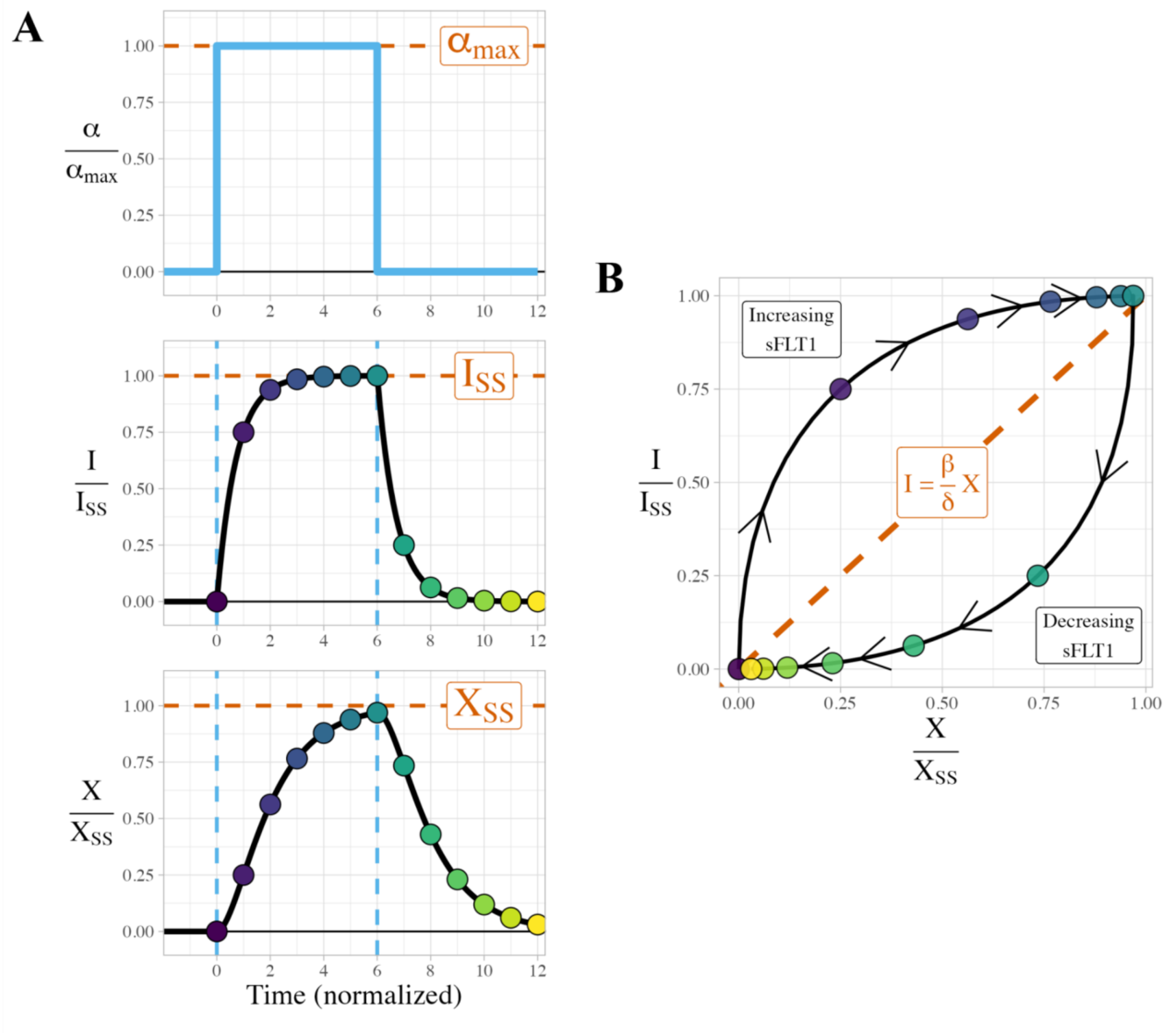


**Supplemental Figure S1. Dynamics and phase plane behavior of an example ODE model solution. (A)** Solution of the ODE model as a function of time showing intracellular ($I$) and extracellular ($X$) sFLT1 protein dynamics in response to step changes in sFLT1 production ($\alpha$). Time is normalized to $T_{50\_X}$, the half-life of $X$ (Eq. 4). $I, X$ are normalized to their steady state values $I_{SS}, X_{SS}$ (Eq. 1-2). **(B)** Phase portrait of the shown ODE model solution. Points are evenly spaced every time unit and colored by time as in panel A. Arrowheads indicate the direction of increasing time starting with $t=0$ in the lower left. The dashed identity line represents potential steady states ($I = \frac{\beta}{\delta} \cdot X$); total sFLT1 increases in regions above this line and decreases below. Parameter values: $\alpha$=10,000, $\beta$=0.1, $\gamma$=0.1, $\delta$= 0.1. For discussion of this figure, see *Supplemental Results*.

Supplemental Figure S2

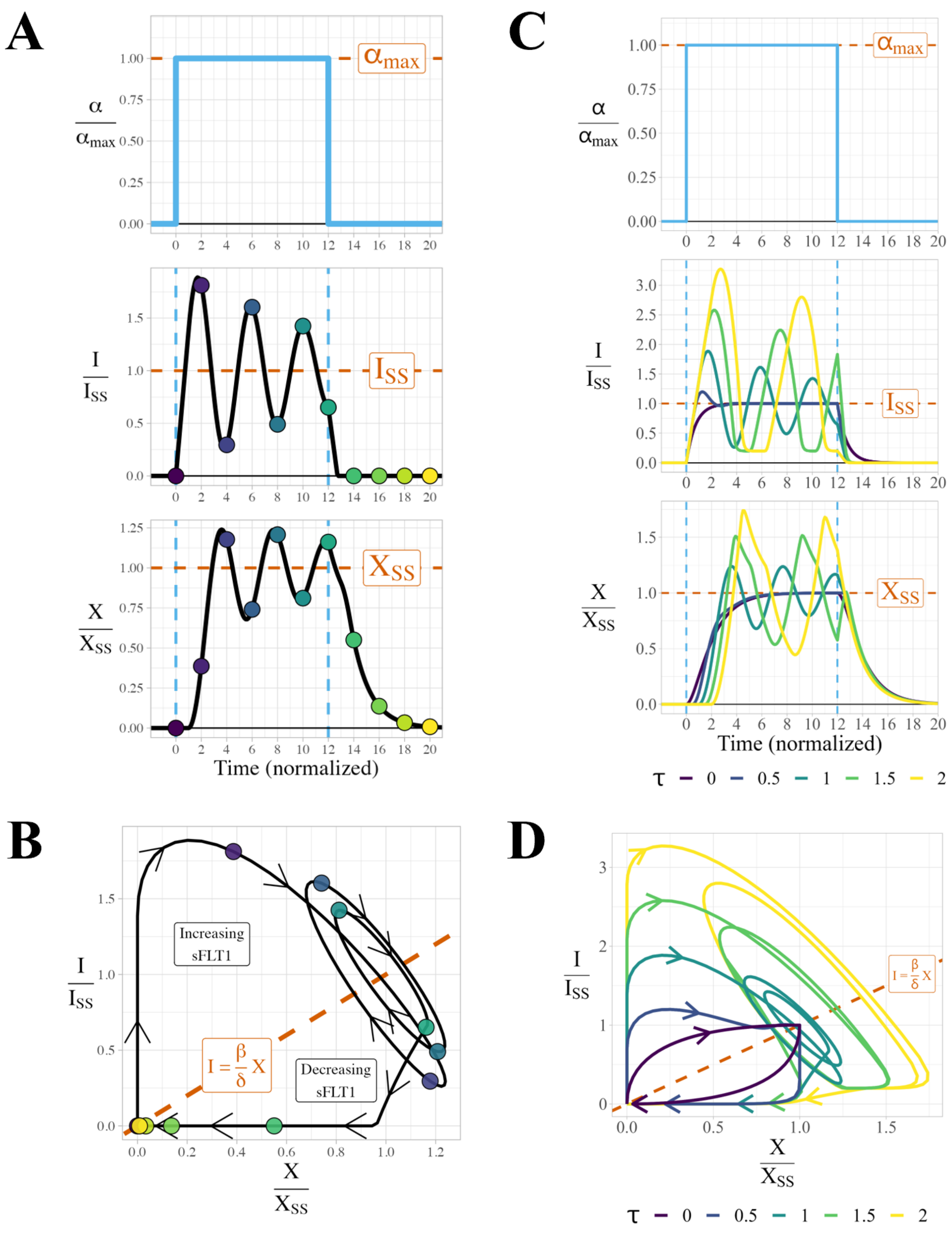


**Supplemental Figure S2. Dynamics and phase plane behavior of an example DDE model solution. (A)** Solution of the DDE model as a function of time showing intracellular (*I*) and extracellular (*X*) sFLT1 protein dynamics in response to step changes in sFLT1 production (α). Time is normalized to $T_{50\_X}$, the half-life of $X$(Eq. 4). $I, X$ are normalized to their steady state values $I_{SS}, X_{SS}$ (Eq. 1-2). **(B)** Phase portrait of the shown DDE model solution. Points are evenly spaced every two time units and colored by time as in panel A. Arrowheads indicate the direction of increasing time starting with t=0 in the lower left. The dashed identity line represents potential steady states $\left(I = \frac{\beta}{\delta} \cdot X\right)$; total sFLT1 increases in regions above this line and decreases below. Parameter values: $\alpha = 10{,}000$, $\beta = 0.1$, $\gamma = 0.1$, $\delta = 0.1$, $\tau = T_{50\_X}$. **(C)** Example DDE solutions as a function of time and **(D)** as a phase portrait for maturation delays $\tau = (0, 0.5, 1, 1.5, 2) \cdot T_{50\_X}$. For discussion of this figure, see *Supplemental Results*.

### Supplemental Figure S3

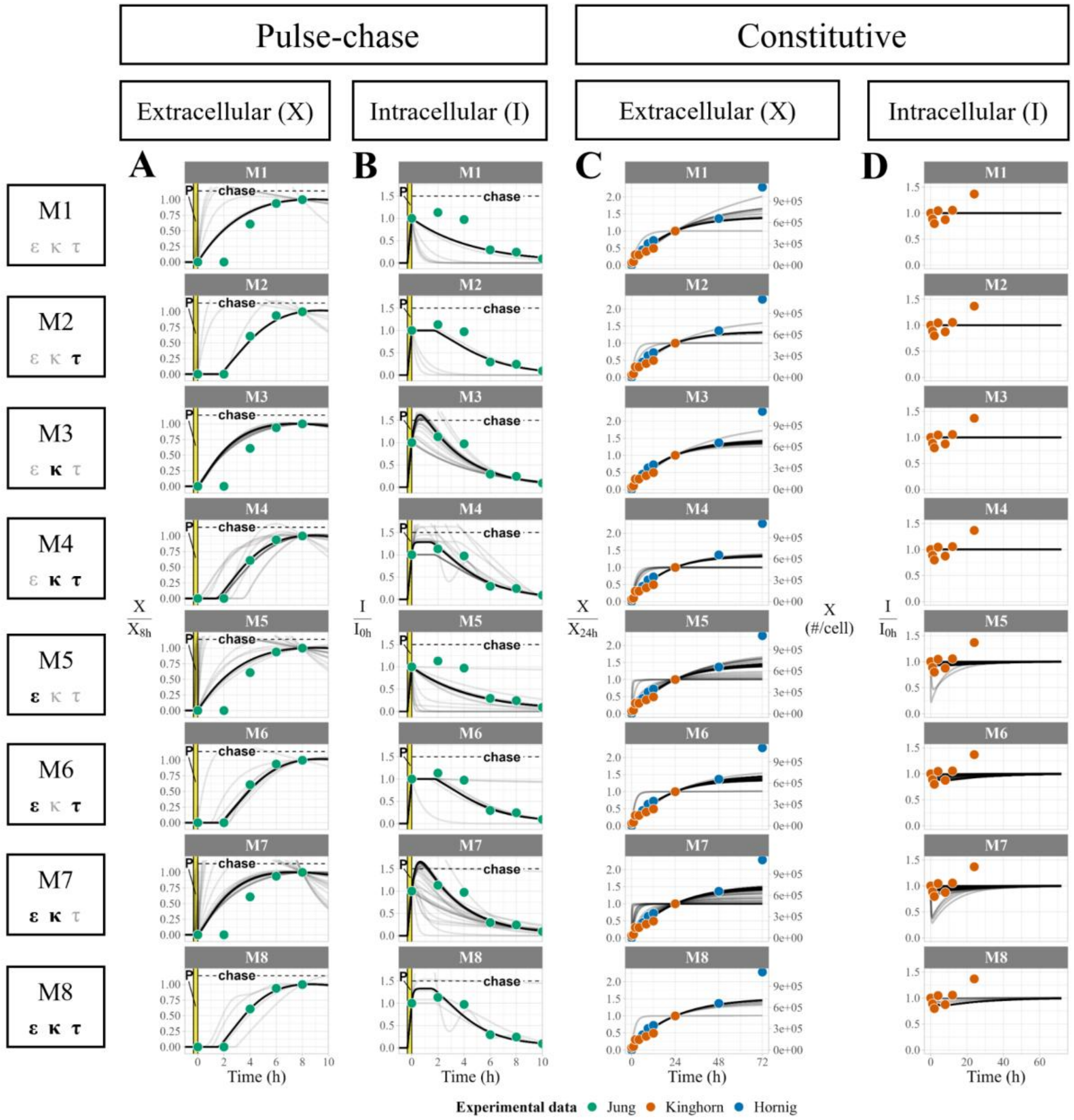


**Supplemental Figure S3. Visual predictive checks of candidate models of sFLT1 secretion.** Comparison of experimental data (points) and simulated time courses (lines, $n = 100$ per plot) of **(A, C)** extracellular and **(B, D)** intracellular sFLT1 for **(A, B)** pulse-chase secretion and **(C, D)** constitutive secretion cases using eight distinct candidate models as described in Figure 3A and Supplemental Table 1. In pulse-chase plots, the highlighted region (P) marks the 20-minute pulse. $X/X_{8h}$: extracellular sFLT1 normalized to its value at 8h; $X/X_{24h}$: extracellular sFLT1 normalized to its value at 24h; $I/I_{0h}$: intracellular sFLT1 normalized to its value at $t = 0$.

Supplemental Figure S4

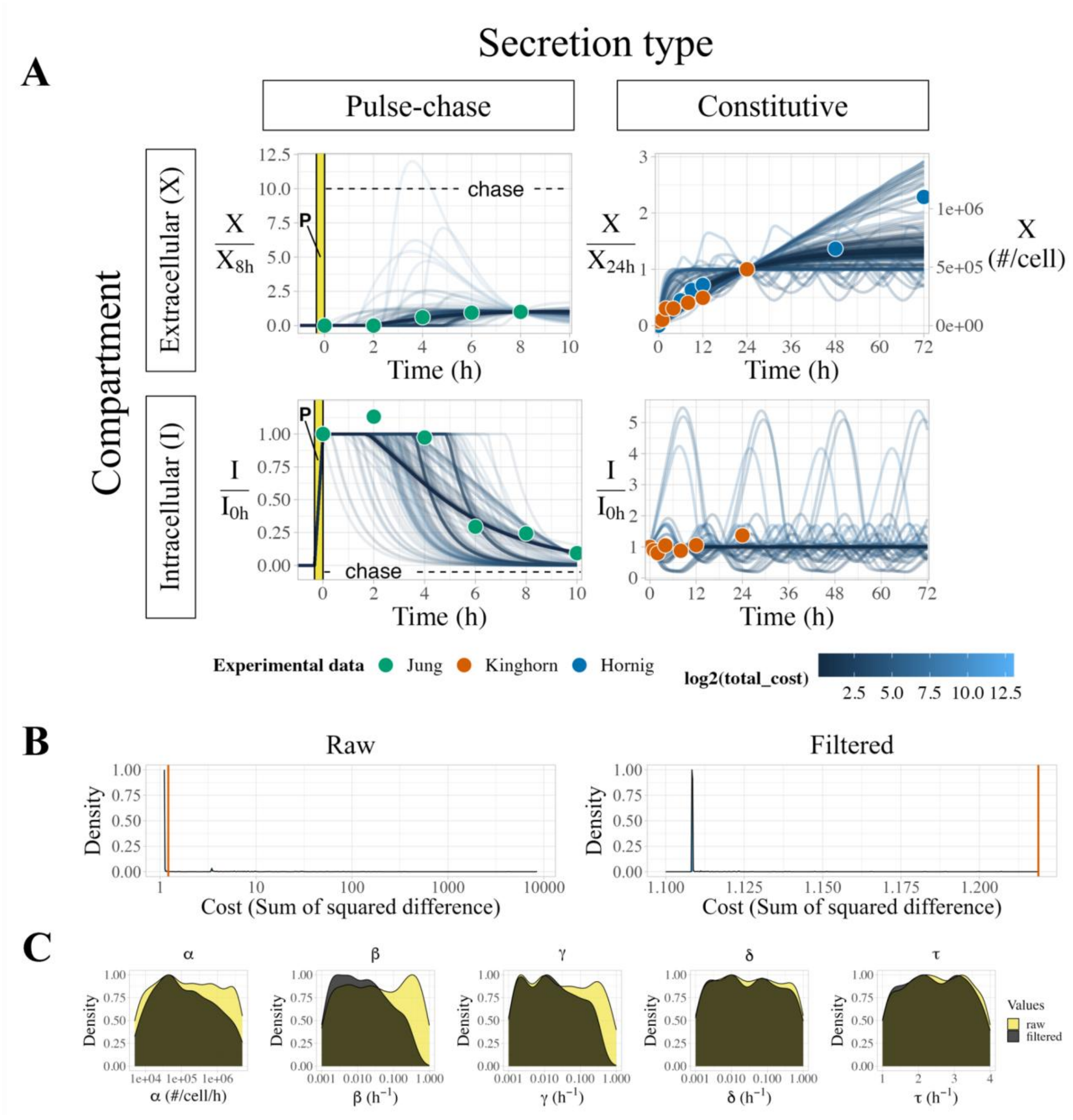


**Supplemental Figure S4. Filtering DDE optimization runs by cost. (A)** Visual predictive checks of unfiltered optimization runs ($n = 1000$) from the delay differential equation (DDE) model. Many runs with high cost have not converged to follow the experimental data. **(B)** Cost distribution of raw (left) and filtered (right) optimization runs of the DDE model. The red lines indicate the cost cutoff of 10% above the minimum cost. **(C)** Density plots of raw and filtered initial parameter values for the DDE model. Raw (yellow) distributions include all attempted optimizations ($n = 1000$). Filtered (black) distributions include only fits with a cost within 10% of the best fit ($n = 594$). Y axes are normalized such that each distribution has a maximum density of 1.

#### Supplemental Figure S5

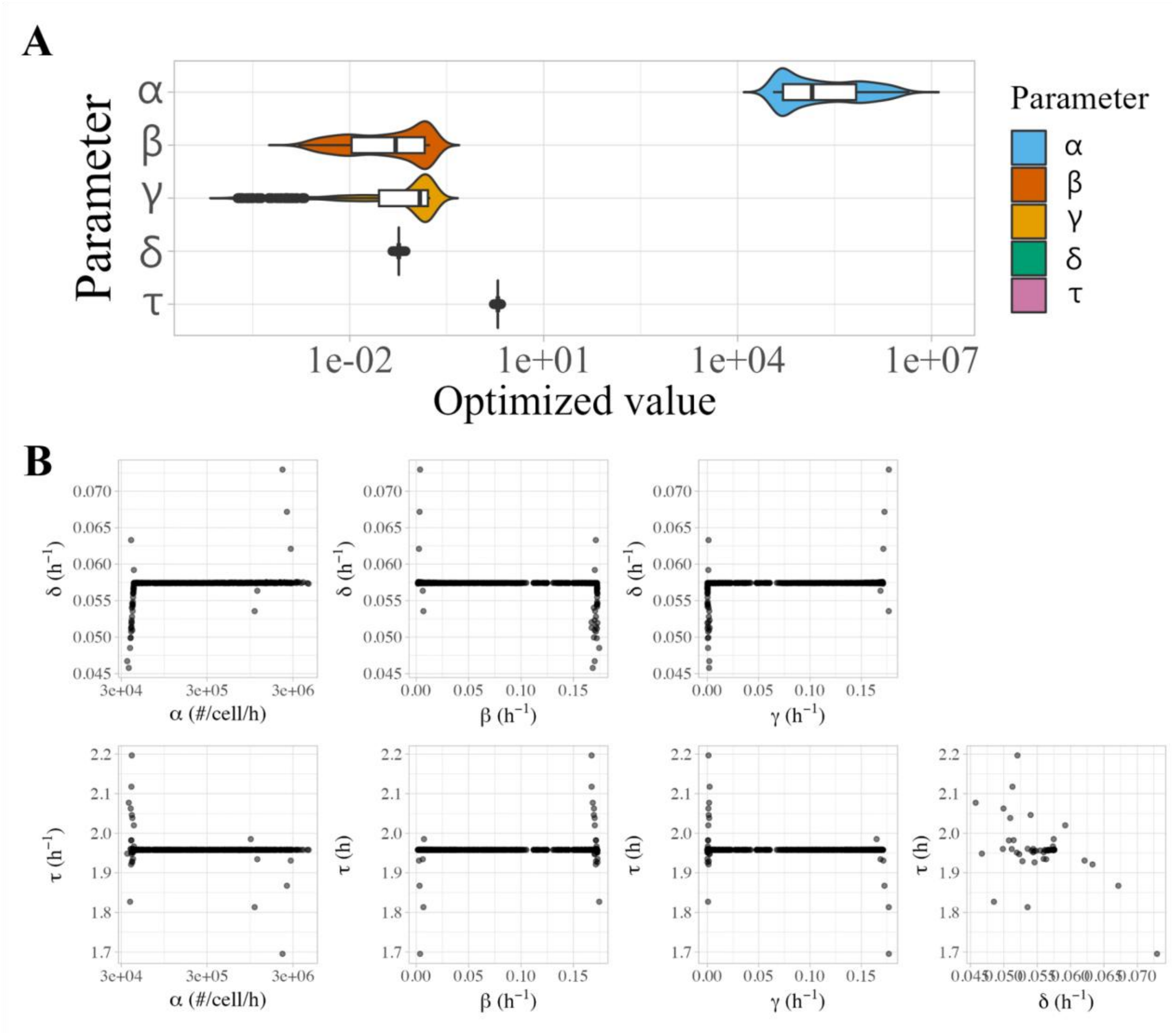


**Supplemental Figure S5. Properties of optimized parameters from the DDE model of sFLT1 secretion. (A)** Violin plots of optimal parameter values ($n = 594$) for the delay differential equation (DDE) model. Units: $\alpha$, #/cell/h; ($\beta$,$\gamma$,$\delta$,$\varepsilon$), $h^{-1}$. **(B)** Correlations between δ and τ and other optimized parameters. Each point represents the observed values of the listed parameters in a single run of the delay differential equation model after filtering for low-cost fits ($n = 594$).

### Supplemental Figure S6

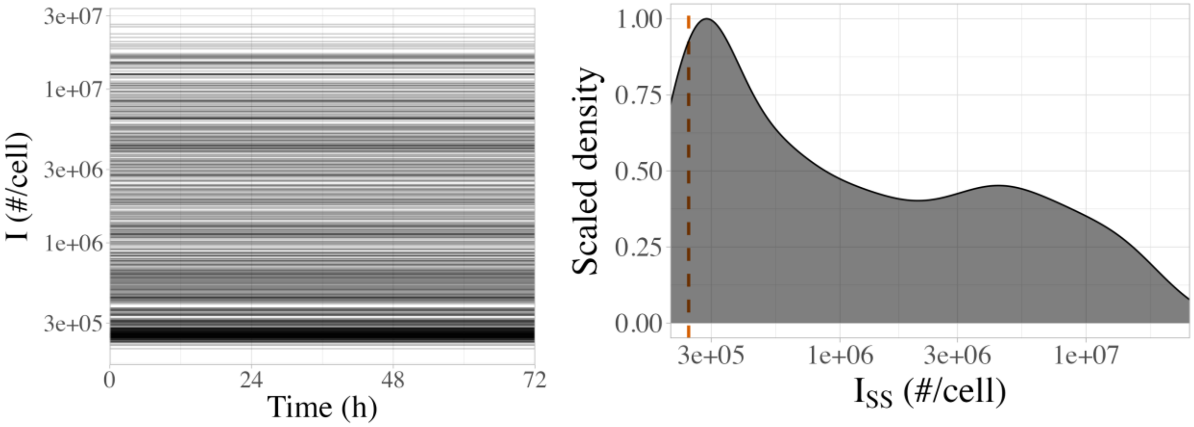


**Supplemental Figure S6. Simulated absolute numbers of intracellular sFLT1 molecules. (A)** Time courses of absolute intracellular sFLT1 ($I$) during simulation of constitutive secretion. **(B)** Distribution of steady state values of intracellular sFLT1 ($I_{SS}$). The dashed red line indicates the theoretical lower bound $\min(I_{SS}) = c_1/c_2^2$ based on median values of $c_1$ and $c_2$ (Table 2).

### Supplemental Figure S7

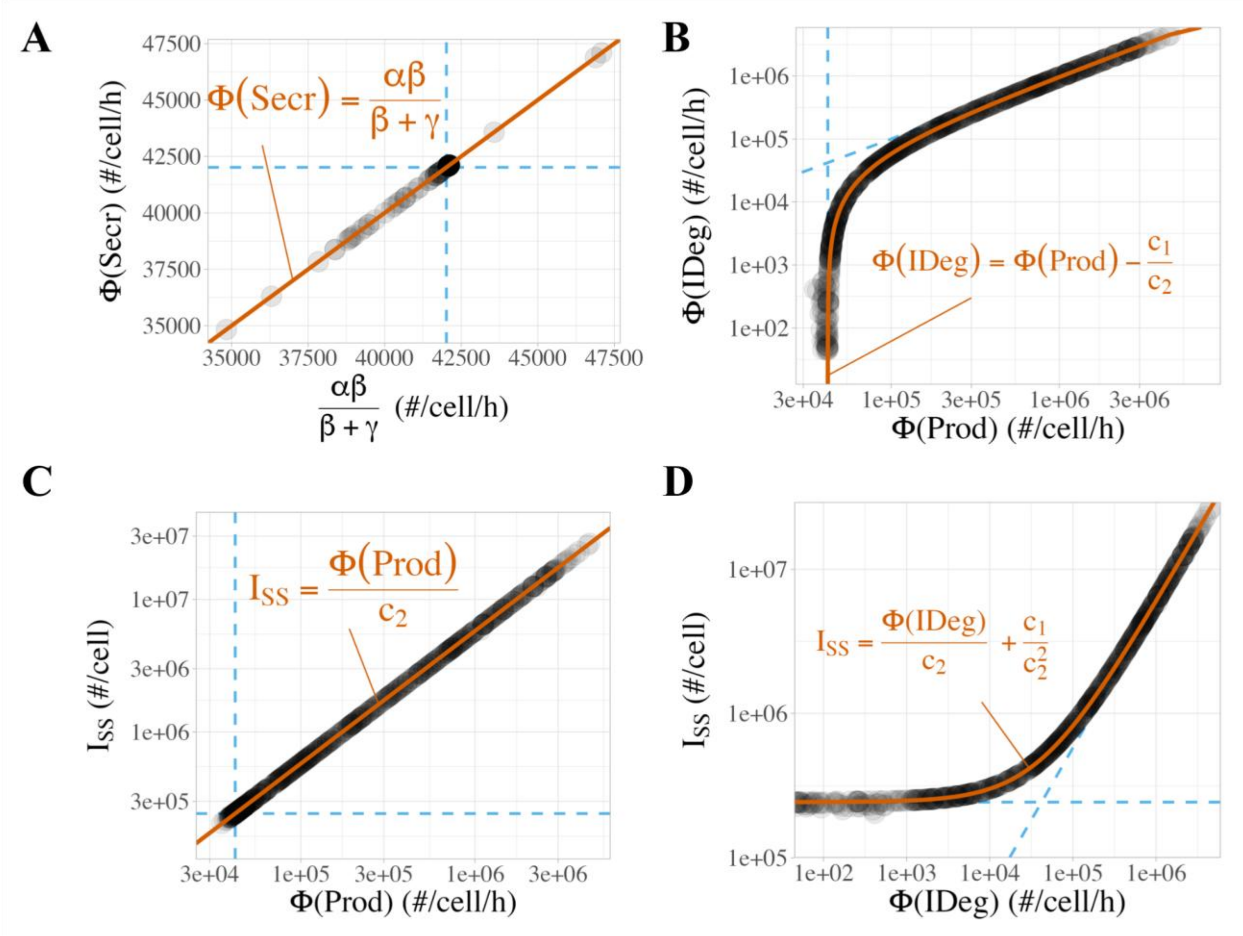


**Supplemental Figure S7. Steady state properties of process fluxes during simulation of constitutive secretion.** All annotation lines are derived in Supplemental Methods and calculated using median values of $c_1$ and $c_2$. **(A)** Correlation between secretion flux (Φ(Secr)) and the compound parameter $\alpha\beta/(\beta+\gamma) = c_1/c_2$ . Both dashed lines indicate $c_1/c_2$ . **(B)** Correlation between intracellular degradation flux (Φ(IDeg)) and production flux (Φ(Prod)). Dashed lines indicate the theoretical lower bound min(Prod) $= c_1/c_2$ and the asymptote Φ(IDeg) = Φ(Prod). **(C)** Correlation between steady state intracellular sFLT1 ($I_{SS}$) and intracellular degradation flux (Φ(IDeg)). Dashed lines indicate the theoretical lower bound $\min(I_{SS}) = c_1/c_2^2$ and the asymptote $I_{SS}$ = Φ(IDeg)/$c_2$. **(D)** Correlation between steady state intracellular sFLT1 ($I_{SS}$) and production flux (Φ(Prod)) during constitutive secretion. Dashed lines indicate theoretical lower bounds ($\min(I_{SS}) = c_1/c_2^2$) and min(Φ(Prod)) $= c_1/c_2$). Φ = Flux, Prod = production, Secr = secretion, IDeg = intracellular degradation, XDeg = extracellular degradation.

Supplemental Figure S8

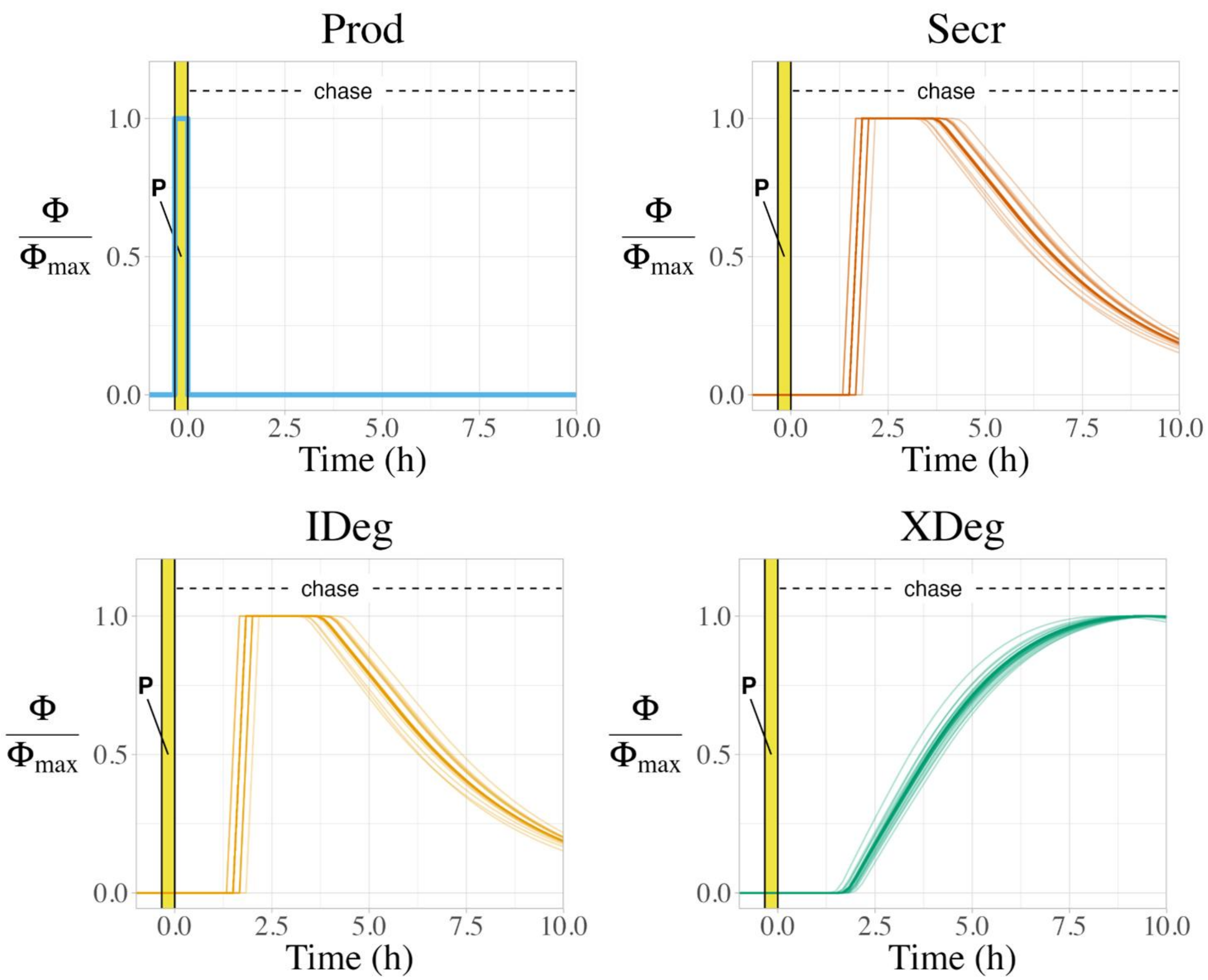


**Supplemental Figure S8. Relative process fluxes over time during simulations of pulse-chase secretion.** All species are normalized to their maximum observed values over 10 hours. Φ = flux, Prod = production, Secr = secretion, IDeg = intracellular degradation, XDeg = extracellular degradation.

### Supplemental Figure S9

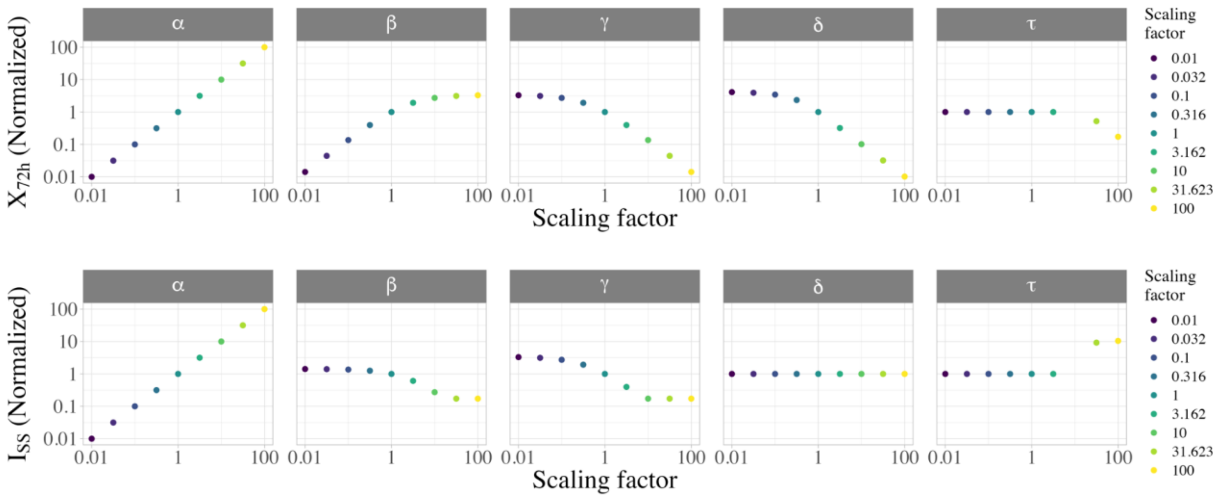


**Supplemental Figure S9. Impact of individual parameter variations on sFLT1 for simulations of constitutive secretion.** Top: extracellular sFLT1 at 72 hours ($X_{72h}$); bottom: steady state intracellular sFLT1 ($I_{SS}$).

### Supplemental Figure S10

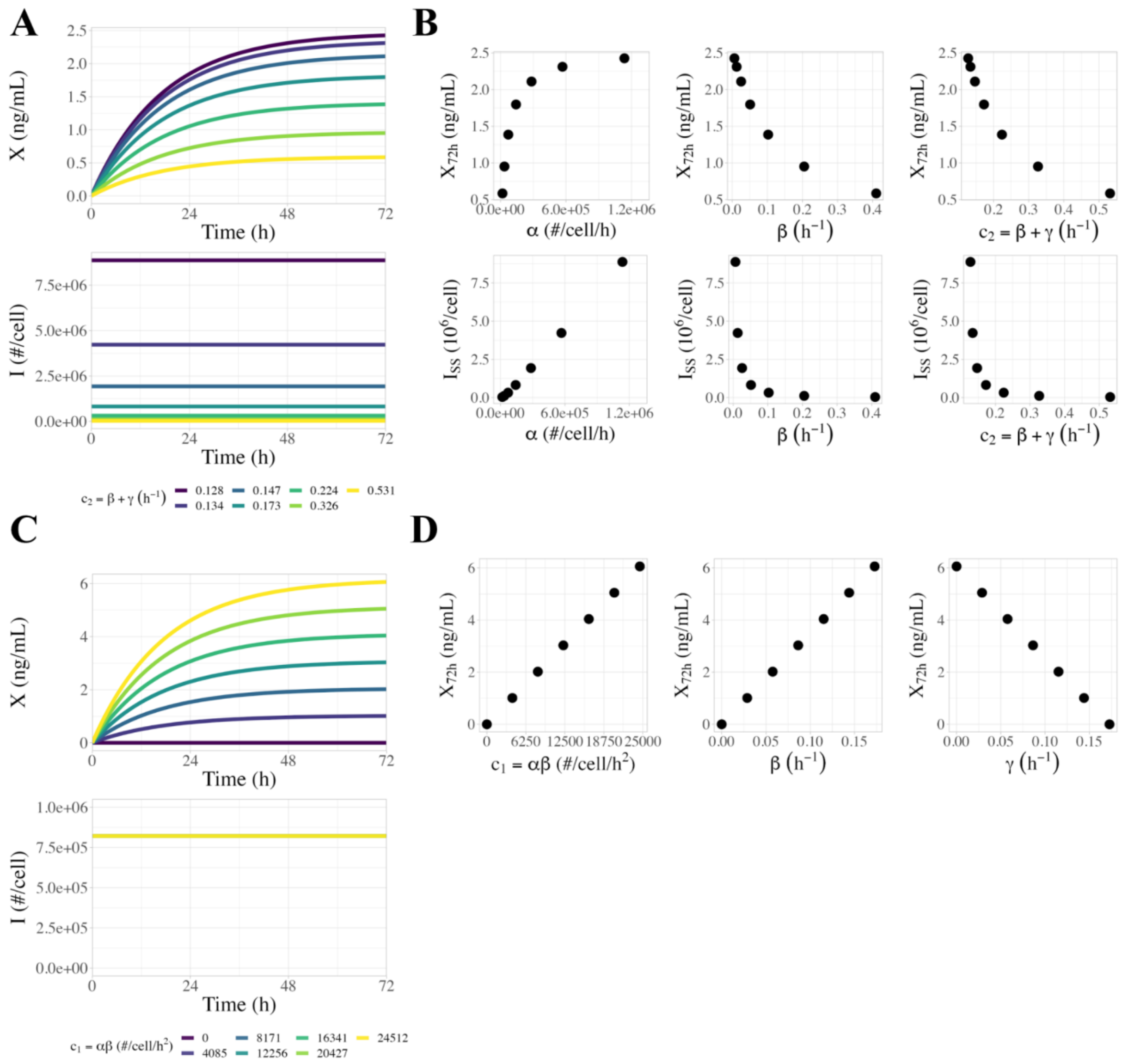


**Figure S10. Sensitivity of extracellular and intracellular sFLT1 to the compound parameters $c_1 = \alpha\beta$ and $c_2 = \beta + \gamma$ in the DDE model. (A)** Time courses of extracellular ($X$) and intracellular ($I$) sFLT1 during constitutive secretion with fixed $c_1 = 7270$ #/cell/h$^2$ and varying $c_2$. **(B)** Correlation between extracellular sFLT1 at 72 hours ($X_{72h}$) or steady state intracellular sFLT1 ($I_{SS}$) with $\alpha$, $\beta$, and $c_2$ when $c_1$ is constant during constitutive secretion. **(C)** Time courses of extracellular (X) and intracellular (I) sFLT1 during constitutive secretion with fixed $c_2 = 0.173$ h$^{-1}$ and varying $c_1$. **(D)** Correlation between extracellular sFLT1 at 72 hours ($X_{72h}$) with $c_1$, $\beta$, and $\gamma$ when $c_2$ is constant during constitutive secretion.

Supplemental Figure S11

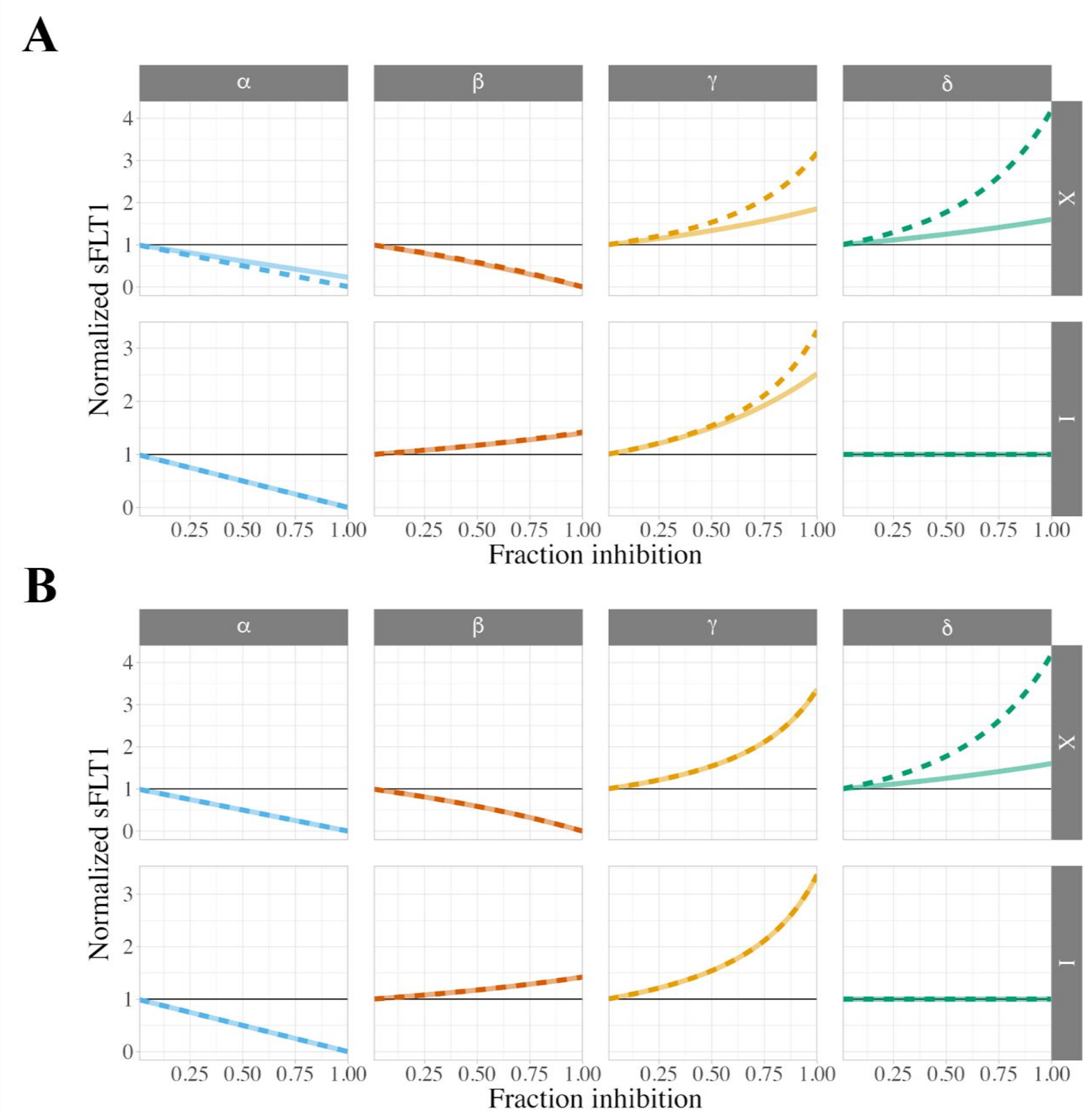


**Supplemental Figure S11. Simulating different strengths of chemical inhibition and genetic downregulation of sFLT1 secretion.** Change in extracellular (X) and intracellular (I) sFLT1 at 18 hours (solid) or 72 hours (dashed) with **(A)** chemical inhibition or **(B)** genetic downregulation of individual parameters at varying fraction inhibition. sFLT1 values are normalized to the 18-hour (solid) or 72-hour (dashed) values from simulation with the median parameter set.

### Supplemental Figure S12

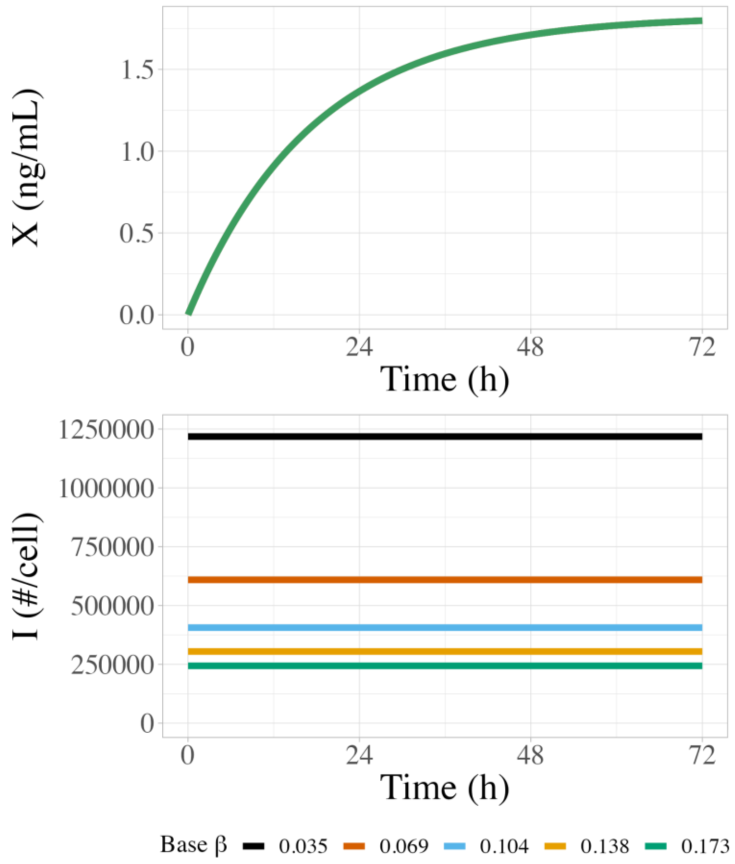


**Supplemental Figure S12. Different sets of initial conditions for simulating chemical and genetic inhibition.** Time courses of extracellular ($X$) and intracellular ($I$) sFLT1 during constitutive simulation with different base parameter sets sharing constraints of $c_1 = \alpha\beta = 7270$ #/cell/h$^2$ and $c_2 = \beta + \gamma = 0.173$ h$^{-1}$. These time courses are used as the base cases for testing chemical and genetic inhibition from different initial conditions (Figure 9, Supplemental Figure S12).

### Supplemental Figure S13

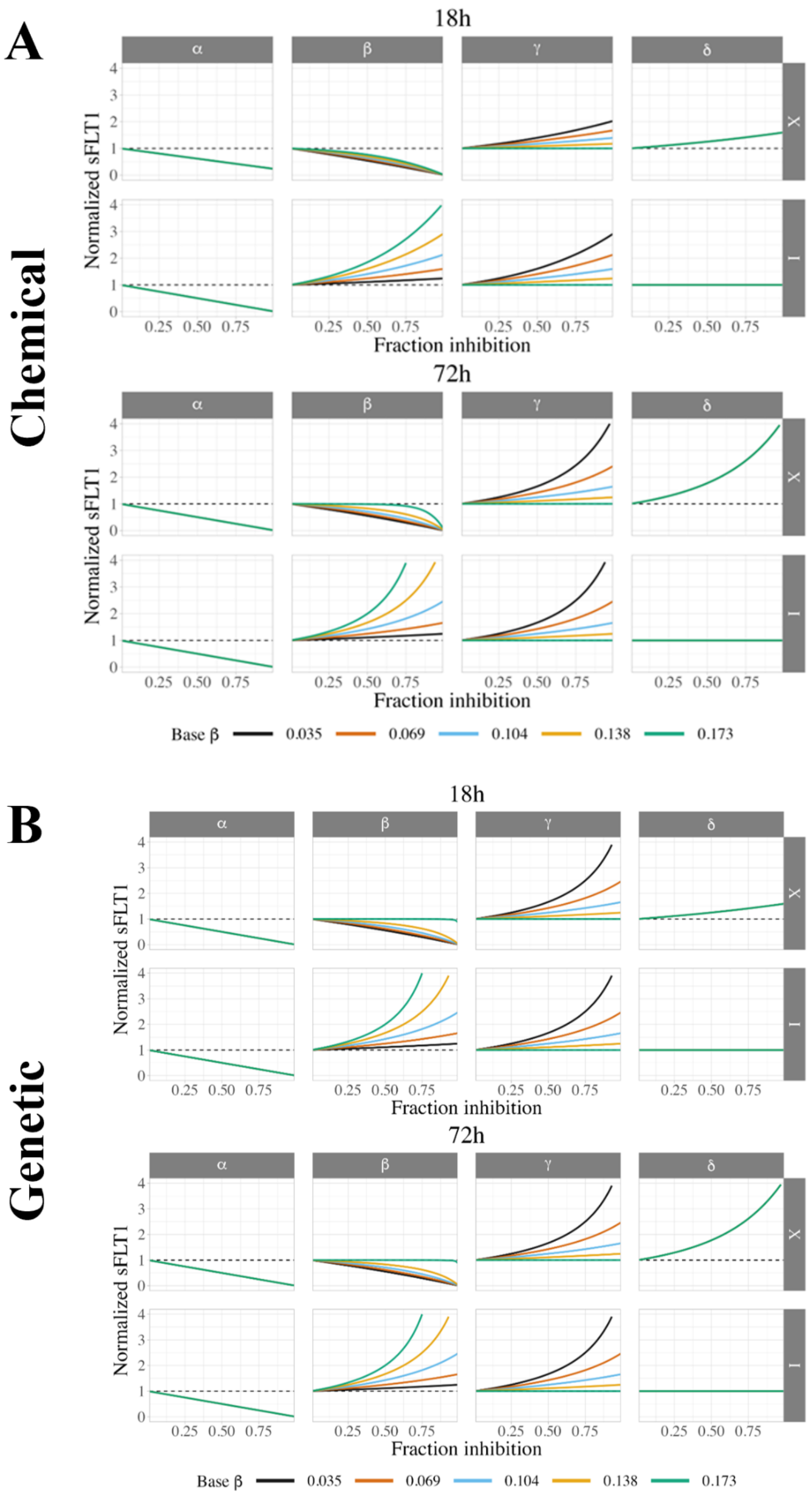


**Supplemental Figure S13. Variation in predicted effects of chemical and genetic perturbations across optimized parameter sets. (A)** Change in extracellular (*X*) and intracellular (*I*) sFLT1 at 18 hours (top) and 72

hours (bottom) with chemical inhibition of individual parameters at varying fraction inhibition for different base parameter sets (labeled by β values). **(B)** Change in extracellular (*X*) and intracellular (*I*) sFLT1 at 18 hours (top) and 72 hours (bottom) post media change with genetic inhibition of individual parameters at varying fraction inhibition for different base parameter sets (labeled by β values). *X* and *I* are normalized to the values at the end of each time course (18h or 72h, respectively) from the simulation with the corresponding base parameter set.

### Supplemental References

**MATLAB software.** MATLAB and Optimization Toolbox Release 2022a, The MathWorks, Inc., Natick, Massachusetts, United States.

**R software.** References were generated in R using `report::cite_packages`.

Bengtsson H (2022). R.matlab: Read and Write MAT Files and Call MATLAB from Within R. R package version 3.7.0, https://CRAN.R-project.org/package=R.matlab.

Garnier, Simon, Ross, Noam, Rudis, Robert, Camargo, Pedro A, Sciaini, Marco, Scherer, Cédric (2023). viridis(Lite) - Colorblind-Friendly Color Maps for R. doi:10.5281/zenodo.4678327 https://doi.org/10.5281/zenodo.4678327, viridisLite package version 0.4.2, https://sjmgarnier.github.io/viridis/.

Garnier, Simon, Ross, Noam, Rudis, Robert, Camargo, Pedro A, Sciaini, Marco, Scherer, Cédric (2024). viridis(Lite) - Colorblind-Friendly Color Maps for R. doi:10.5281/zenodo.4679423 https://doi.org/10.5281/zenodo.4679423, viridis package version 0.6.5, https://sjmgarnier.github.io/viridis/.

Grolemund G, Wickham H (2011). "Dates and Times Made Easy with lubridate." Journal of Statistical Software, 40(3), 1-25. https://www.jstatsoft.org/v40/i03/.

Gu Z, Eils R, Schlesner M (2016). "Complex heatmaps reveal patterns and correlations in multidimensional genomic data." Bioinformatics. doi:10.1093/bioinformatics/btw313 https://doi.org/10.1093/bioinformatics/btw313. Gu Z (2022). “Complex Heatmap Visualization.” iMeta. doi:10.1002/imt2.43 https://doi.org/10.1002/imt2.43.

Gu Z, Gu L, Eils R, Schlesner M, Brors B (2014). “circlize implements and enhances circular visualization in R.” Bioinformatics, 30, 2811-2812.

Meschiari S (2022). latex2exp: Use LaTeX Expressions in Plots. R package version 0.9.6, https://CRAN.R-project.org/package=latex2exp.

Müller K, Wickham H (2023). tibble: Simple Data Frames. R package version 3.2.1, https://CRAN.R-project.org/package=tibble.

Pedersen T (2024). patchwork: The Composer of Plots. R package version 1.2.0, https://CRAN.R-project.org/package=patchwork.

R Core Team (2024). R: A Language and Environment for Statistical Computing. R Foundation for Statistical Computing, Vienna, Austria. https://www.R-project.org/.

Schloerke B, Cook D, Larmarange J, Briatte F, Marbach M, Thoen E, Elberg A, Crowley J (2024). GGally: Extension to 'ggplot2'. R package version 2.2.1, https://CRAN.R-project.org/package=GGally.

Slowikowski K (2024). ggrepel: Automatically Position Non-Overlapping Text Labels with 'ggplot2'. R package version 0.9.6, https://CRAN.R-project.org/package=ggrepel.

Wickham H (2011). “testthat: Get Started with Testing.” The R Journal, 3, 5-10. https://journal.r-project.org/archive/2011-1/RJournal_2011-1_Wickham.pdf.

Wickham H (2016). ggplot2: Elegant Graphics for Data Analysis. Springer-Verlag New York. ISBN 978-3-319-24277-4, https://ggplot2.tidyverse.org.

Wickham H (2023). forcats: Tools for Working with Categorical Variables (Factors). R package version 1.0.0, https://CRAN.R-project.org/package=forcats.

Wickham H (2023). stringr: Simple, Consistent Wrappers for Common String Operations. R package version 1.5.1, https://CRAN.R-project.org/package=stringr.

Wickham H, Averick M, Bryan J, Chang W, McGowan LD, François R, Grolemund G, Hayes A, Henry L, Hester J, Kuhn M, Pedersen TL, Miller E, Bache SM, Müller K, Ooms J, Robinson D, Seidel DP, Spinu V, Takahashi K, Vaughan D, Wilke C, Woo K, Yutani H (2019). “Welcome to the tidyverse.” Journal of Open Source Software, 4(43), 1686. doi:10.21105/joss.01686 https://doi.org/10.21105/joss.01686.

Wickham H, François R, Henry L, Müller K, Vaughan D (2023). dplyr: A Grammar of Data Manipulation. R package version 1.1.4, https://CRAN.R-project.org/package=dplyr.

Wickham H, Henry L (2023). purrr: Functional Programming Tools. R package version 1.0.2, https://CRAN.R-project.org/package=purrr.

Wickham H, Hester J, Bryan J (2024). readr: Read Rectangular Text Data. R package version 2.1.5, https://CRAN.R-project.org/package=readr.

Wickham H, Pedersen T, Seidel D (2023). scales: Scale Functions for Visualization. R package version 1.3.0, https://CRAN.R-project.org/package=scales.

Wickham H, Vaughan D, Girlich M (2024). tidyr: Tidy Messy Data. R package version 1.3.1, https://CRAN.R-project.org/package=tidyr.

Wilke C, Wiernik B (2022). gridtext: Improved Text Rendering Support for 'Grid' Graphics. R package version 0.1.5, https://CRAN.R-project.org/package=gridtext.
